# Modulation of lipid metabolism by thapsigargin inhibits hepatitis C virus infection by multiple mechanisms

**DOI:** 10.64898/2026.03.02.709032

**Authors:** Trinity H. Tooley, Abbey J. McMurray, Isabella E. Pellizzari-Delano, Sophia K. Stegeman, Katrina Gee, Che C. Colpitts

## Abstract

Thapsigargin (Tg), an inducer of endoplasmic reticulum stress and the unfolded protein response (UPR), has broad-spectrum antiviral activity, although the underlying mechanisms remain unclear. Here, we characterized its antiviral mechanism(s) using hepatitis C virus (HCV) as a model. Pre-treatment with Tg partially inhibited HCV RNA replication, but more strongly reduced extracellular viral titer and RNA, suggesting a block to later stages of infection. Its antiviral effects against HCV infection are likely UPR-independent, since treatment with tunicamycin, which activates the UPR by a different mechanism, did not exert the same antiviral effect. Given the importance of lipid droplets (LDs) and lipid metabolism in mediating HCV assembly and egress, we examined Tg-mediated effects on lipid homeostasis. Tg treatment upregulated the expression of lipid synthesis genes, including *FASN* and *DGAT1/2*, and led to the accumulation of enlarged LDs. Tg also induced expression of *CIDE-C*, a mediator of LD fusion. Silencing CIDEC expression impaired Tg-induced LD enlargement and rescued viral RNA replication, but not extracellular titer, from Tg inhibition, demonstrating that Tg-mediated LD remodeling contributes to replication defects without significantly affecting assembly or egress. Intracellular viral titers were unchanged in Tg-treated cells, indicating intact virion assembly but a defect in secretion. Consistently, Tg treatment reduced both apolipoprotein B and E secretion, but not that of Gaussia luciferase, suggesting that specific disruption of the very low-density lipoprotein secretion pathway by Tg could impair efficient HCV egress. Together, our findings reveal that modulation of lipid homeostasis by Tg inhibits HCV RNA replication and egress by distinct mechanisms. This work has antiviral implications for other viruses that rely on lipid metabolism during infection.

**AUTHOR’S SUMMARY:** Thapsigargin (Tg) is emerging as a potent broad-spectrum host-targeted antiviral, but the specific mechanisms by which it inhibits unrelated viral infections are unclear and its antiviral activity against hepatitis C virus has not been reported. We determined that Tg causes significant changes in host lipid metabolism, resulting in enlarged lipid droplets (LDs) that persist during HCV infection. We demonstrate that Tg-induced enlarged LDs in part contribute to the restriction of HCV RNA replication. However, the most compelling antiviral effect is on the release of infectious virions. We report no differences in intracellular titer, but a significant reduction in extracellular titer, concomitant with a decrease in apolipoprotein (Apo) B and ApoE secretion, which are required for efficient egress of HCV. Together, our data support modulation of lipid metabolism and cellular secretory function by Tg as a previously unidentified antiviral mechanism for Tg and highlight its potential use against other viruses.

## INTRODUCTION

Hepatitis C virus (HCV) is a positive-sense single-stranded RNA virus (+ssRNA) belonging to the *Flaviviridae* family. Despite the advent of highly effective direct acting antivirals (DAAs), the WHO estimates that over 51 million people live with chronic HCV infection (1). While the possibility of eradicating HCV infection relies on the development of an effective vaccine, DAAs have helped significantly in curing HCV-infected patients and limiting spread. Unfortunately, routine use of DAAs has led to increased selective pressure leading to viral resistance, which could potentially render DAAs ineffective (2, 3). One possibility to mitigate the threat of viral resistance to current antivirals is the development of host-targeted antivirals (HTAs). By leveraging our understanding of how viruses, such as HCV, interact with host cell factors to support infection, we may be able to exploit these interactions to develop broadly acting HTAs that protect against unrelated viral infections.

One such HTA is a naturally occurring molecule called thapsigargin (Tg), derived from the Mediterranean plant *Thapsia garganica.* Tg is well-established to induce endoplasmic reticulum (ER) stress by irreversibly binding to the sarcoplasmic/endoplasmic reticulum calcium ATPase (SERCA) pump (4), thereby depleting calcium levels within the ER lumen and leading to the activation of the unfolded protein response (UPR). Tg has broad-spectrum antiviral activity against many +ssRNA viruses, including coronaviruses (5, 6) and viruses within the *Flaviviridae* family, such as dengue virus (7), Zika virus (8, 9) and tick-borne encephalitis virus (10). Interestingly, it has been reported that a short 30-minute treatment with Tg in vitro confers a long-lasting antiviral effect against influenza A virus (11). However, its specific antiviral mechanisms are still unclear.

Tg treatment activates the UPR, which is comprised of three distinct signalling pathways that are regulated by three transmembrane ER resident sensors: Activating transcription factor 6 (ATF6), PKR-like endoplasmic reticulum kinase (PERK) and inositol requiring enzyme 1α (IRE1α) (12). The UPR functions to maintain ER homeostasis by transcriptionally upregulating genes to restore proteostasis, halting protein translation through the phosphorylation of eukaryotic initiation factor 2α (eIF2α), or promoting cellular apoptosis if ER homeostasis cannot be restored (12). While it has been speculated that induction of ER stress and UPR activation mediate the antiviral effect of Tg (6), we recently found that the antiviral effect of Tg against the human common cold coronavirus HCoV-229E does not require individual UPR sensor expression (13), suggesting that its antiviral activity is at least partially UPR-independent.

Another consequence of ER stress is the modulation of lipid homeostasis, which can occur through both UPR-dependent and independent mechanisms. For example, activation of the UPR can alter expression of key lipid-related enzymes, thus influencing lipid synthesis (lipogenesis) or breakdown of lipids (lipolysis) (14). ER stress also modulates lipid metabolism through UPR-independent mechanisms, primarily through alterations in ER lipids or calcium depletion, which can promote activation of sterol regulatory element binding proteins (SREBPs). SREBP1 and SREBP2 are central regulators of cellular lipid metabolism and cholesterol biosynthesis, respectively (15). Tg treatment has been shown to increase activation of SREBP2 and accumulation of free cholesterol (16). Ultimately, activation of SREBP2 can lead to an increase in expression of genes that regulate cholesterol and lipoprotein uptake (17), highlighting additional mechanisms through which Tg can modulate lipid metabolism.

Tg treatment upregulates the expression of genes encoding lipogenic trans activators, as well as enzymes involved in lipid droplet (LD) formation and triglyceride synthesis in a human hepatoma cell line (18). Consistent with an increase in expression of lipogenic genes, Tg treatment induces the formation of cytoplasmic LDs at early time points post-treatment (18) and has been reported to also induce LD enlargement (19). LDs are dynamic organelles that store neutral lipids, such as cholesterol esters and triacylglycerides (20), and play integral roles in supporting the replication cycle of many viruses (21). For example, the HCV core (capsid) protein localizes to LDs, which has been proposed to aid in nucleocapsid assembly and promote the production of progeny virions (22–25). Furthermore, HCV egress is closely linked to the lipoprotein secretion pathway, resulting in the production of lipoviral particles that are associated with low-density lipoproteins such as apolipoprotein (Apo) B and E, which allow for more efficient viral entry and evasion from host antibodies (26). LDs are also emerging as regulators of innate antiviral immunity, suggesting another key function at the virus-host interface (27, 28). As such, we hypothesized that Tg-induced alterations to lipid metabolism and LDs could have antiviral consequences for many +ssRNA viruses, including HCV.

In this study, we sought to determine the role of lipid metabolism in the antiviral activity of Tg. We use HCV as a model given the well-characterized role of LDs and lipoprotein secretion in HCV infection. We show that pre-treatment with Tg inhibits HCV infection with the most prominent effect on viral titer, suggesting a disruption to late stages of the viral life cycle. Tg treatment upregulated the expression of LD-related genes and increased the average size of LDs. Silencing expression of cell death inducing DFFA-like effector C (CIDE-C), a mediator of LD fusion, prevented Tg-induced LD enlargement and rescued RNA replication, but not viral extracellular titer, from Tg inhibition. Quantification of intracellular viral titer revealed no difference in DMSO or Tg-treated cells, indicating that Tg disrupts HCV secretion. Tg treatment decreased ApoB and ApoE secretion, reflecting disruption to the host lipoprotein secretion pathway, which is required for efficient HCV egress. Overall, our findings highlight distinct mechanisms through which Tg modulation of host lipid metabolism can impair viral infection.

## RESULTS

### Tg inhibits HCV infection at a late stage

We first established that Tg treatment inhibits HCV infection. We and others have reported that a short 30-minute exposure at low concentrations of Tg is sufficient to cause an antiviral effect (11, 13). As such, we confirmed that a 30-minute pre-treatment with 100 nM Tg broadly induced ER stress, as measured by UPR pathway activation **(Figure 1A)**. We next examined if Tg treatment inhibits HCV infection (HCVcc) by assessing viral titer. Huh7.5 hepatoma cells were pre-treated with DMSO or Tg (100 nM) for 30 minutes and subsequently infected with HCV Japanese Fulminant Hepatitis 1 strain (JFH1_T_ (29)) (MOI 0.2) for 72 h. Tg treatment caused a ∼100-fold decrease in viral titer, indicating that Tg is antiviral against HCV infection **(Figure 1B)**. While we observed that Tg treatment at this concentration resulted in a ∼10% decrease in cell viability **(Figure 1C)**, this is not sufficient to explain the ∼2 log reduction in viral titer.

**Figure 1.**
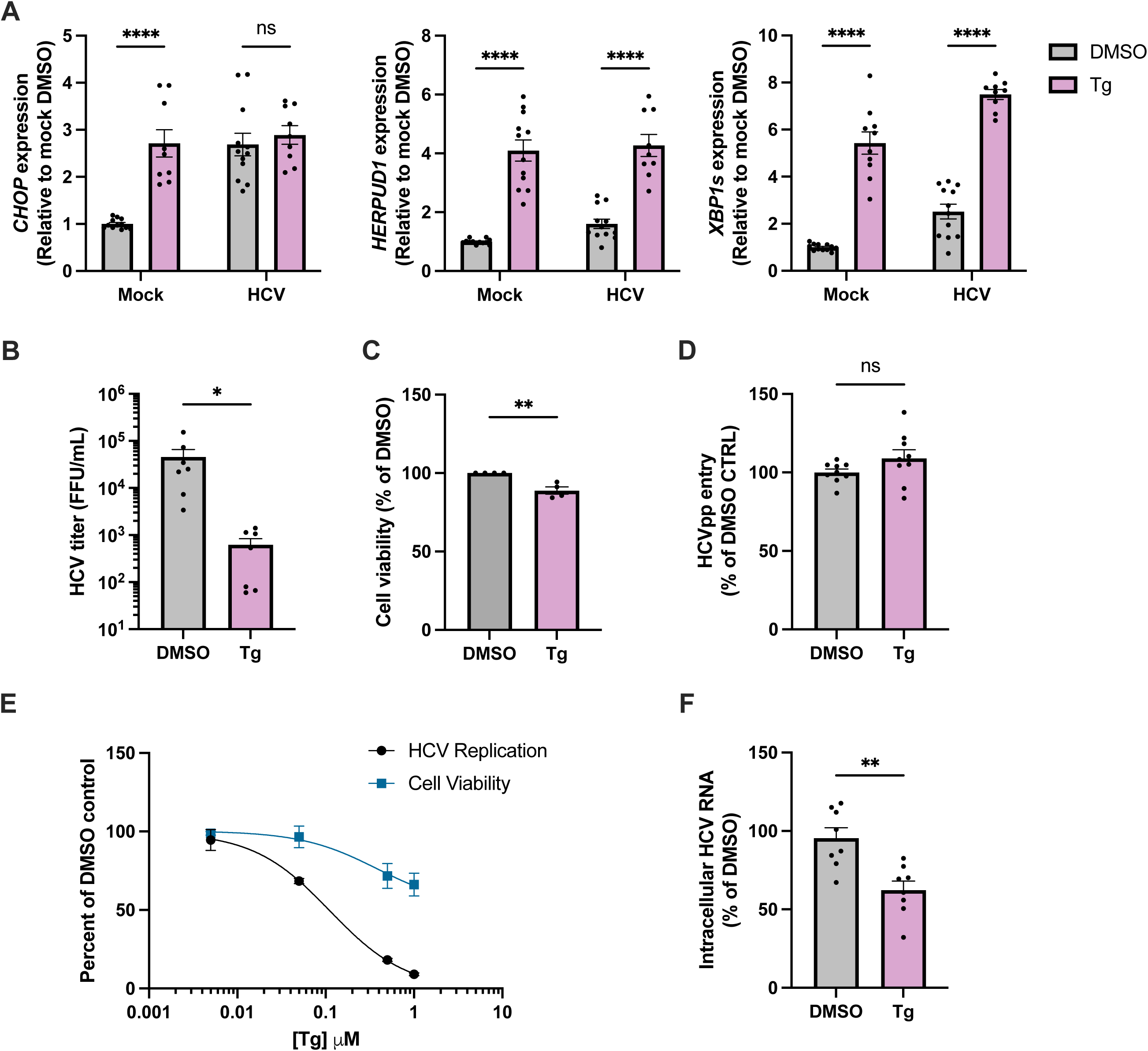
Pre-treatment with ER stress inducer thapsigargin (Tg) reduces HCVcc infection at a post-entry stage. (**A-C**) Huh7.5 cells were pre-treated with DMSO or 100 nM Tg for 30 minutes. **(A)** RNA lysates from mock-infected cells were collected 72 hours post-treatment and UPR gene expression was assessed by RT-qPCR (*HERPUD1*, *CHOP* and *XBP1*). **(B)** DMSO or Tg-treated cells were infected with HCVcc JFH1_T_ (MOI 0.2) for 72 hours. Cellular supernatant was collected and viral titer was assessed by focus-forming assay. **(C)** Huh7.5 cells were treated with DMSO and 100 nM Tg for 30 minutes, and cell viability was assessed 48 hours post treatment by Alamar blue assay. **(D)** DMSO or Tg-treated cells were exposed to HCVpp for 72 hours. Luciferase activity was measured, and data is normalized to DMSO control. **(E)** Huh7 cells were electroporated with 5 µg of HCV SGR RNA. At 4 h post-electroporation, cells were treated with DMSO or various concentrations of Tg for 30 minutes. RNA replication and cell viability was measured by luciferase activity and Alamar blue assay, respectively, at 72 h post-electroporation. **(F)** Huh7.5 cells were pre-treated with DMSO or 100 nM Tg for 30 minutes and subsequently infected with HCVcc JFH1_T_ (MOI 0.2) for 72 h. **(F)** HCVcc intracellular RNA expression was measured by RT-qPCR. Graphs show means +/- SEM from two to six independent experiments. RT-qPCR and reporter assay performed in technical triplicate per biological replicate. *p<0.05, **p<0.01, ***p<0.0005, ****p<0.0001.

To determine which step of the viral life cycle is inhibited by Tg, we first examined viral entry using HCV pseudoparticles (HCVpp) harbouring a luciferase reporter. Huh7.5 cells were pre-treated with DMSO or Tg for 30 minutes and subsequently exposed to HCVpp for 72 hours. Luciferase activity, a proxy for HCVpp entry, was not affected by Tg treatment, indicating that Tg does not impact viral entry **(Figure 1D)**. To test the effect on HCV RNA replication, we electroporated Huh7 cells with an HCV subgenomic replicon (30) (containing only the non-structural proteins required for HCV replication) and treated the electroporated cells with increasing concentrations of Tg for 30 minutes. HCV RNA replication and cell viability was assessed at 72 h post-treatment. The HCV subgenomic replicon contains a luciferase reporter gene downstream of an internal ribosomal entry site, thus allowing cap-independent luciferase expression to be measured as a proxy for viral replication. We observe that treatment with Tg modestly reduced HCV RNA replication at concentrations that did not impact cell viability **(Figure 1E)**. To confirm that Tg affects HCV RNA replication in the context of infection, we infected Huh7.5 cells with HCVcc as described above and harvested intracellular RNA at 72 hpi. HCVcc-infected cells that were pre-treated with Tg (100 nM) displayed a ∼50% reduction in intracellular HCVcc RNA **(Figure 1F)**. Overall, Tg exhibited the strongest effect on viral titer, suggesting that it inhibits later steps of the viral lifecycle, such as assembly or egress. Based on the known effect of UPR activation on lipid metabolism (31), we hypothesized that Tg treatment alters lipid metabolism, which affects nucleocapsid assembly or particle secretion to explain the decrease in viral titer.

### Tg treatment increases lipogenic gene expression and promotes formation of enlarged LDs

To confirm that treatment with Tg alters lipid metabolism in our model, we performed bulk RNA sequencing to assess expression of relevant genes **(Table S1)**. Huh7.5 cells were treated with DMSO or Tg for 30 minutes and cellular RNA was harvested 48 h post-treatment. Reactome gene set enrichment analysis revealed that there is a significant upregulation of genes related to cholesterol biosynthesis **(Figure 2A),** including acetyl-CoA acyltransferase 2 (*ACAT2)*, 3-hydroxy-3-methylgluteryl-CoA-reductase (*HMGCR*), squalene epoxidase (*SQLE*) and others **(Figure S1A)**, confirming that key genes in lipid metabolism are upregulated by Tg treatment. HCV nucleocapsid assembly is thought to occur at LDs, requiring both core and NS5A to localize to LDs, which occurs in a manner that is largely dependent on diacylglycerol acytransferase 1 (DGAT1) (32). NS5A is proposed to shuttle newly synthesized viral RNA from the replication organelle (RO) to LDs, where it is encapsidated by core (33). ER stress causes alterations in lipid homeostasis, and a previous study showed that Tg treatment leads to enlargement of LDs (19). Thus, given the effect of Tg on viral titer **(Figure 1B)** and the role of LDs in facilitating viral assembly, we investigated the effect of Tg on LDs in the context of HCV infection.

**Figure 2.**
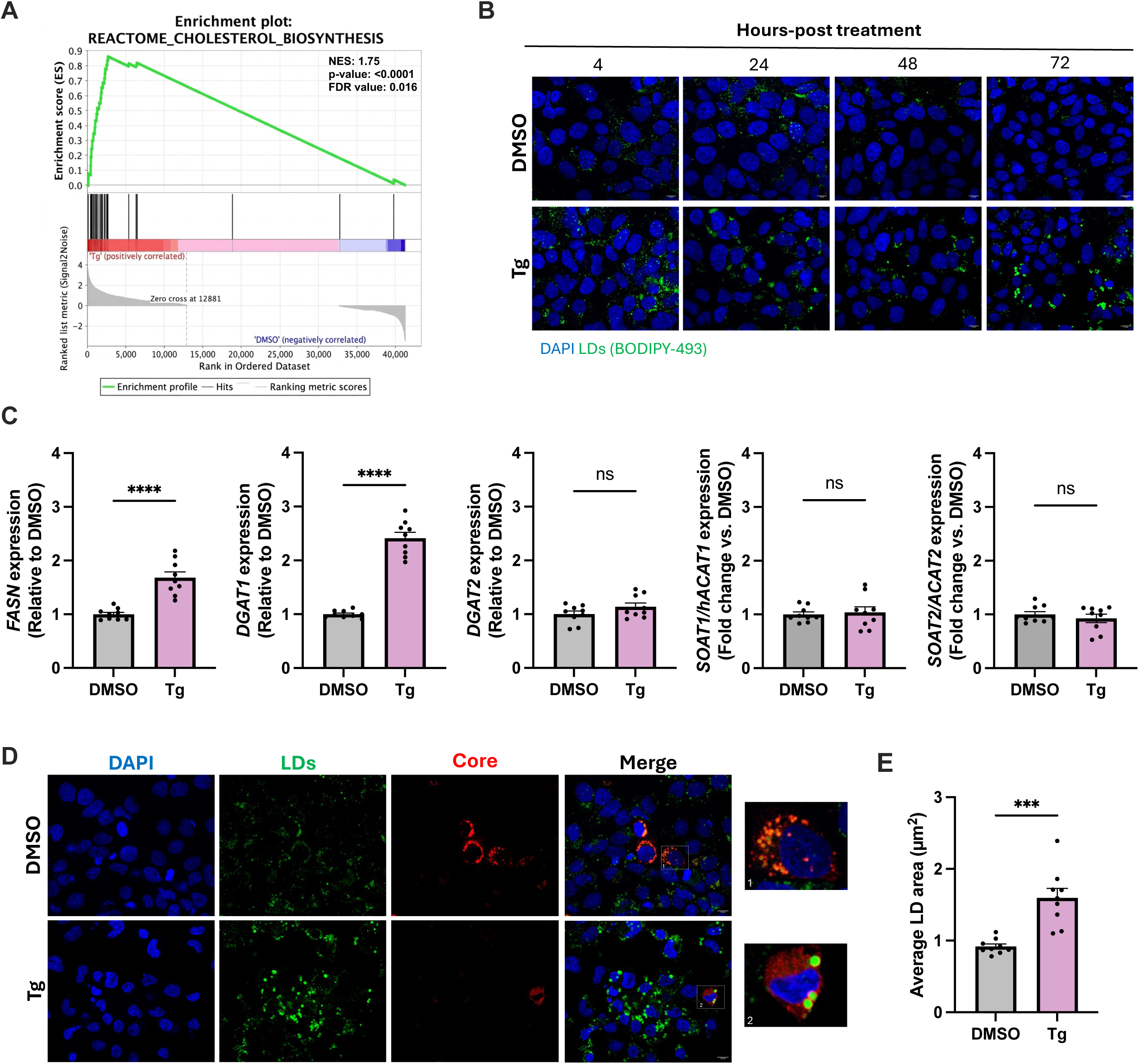
Tg treatment induces lipogenic gene expression and induces the accumulation of enlarged lipid droplets. **(A)** Huh7.5 cells were treated with DMSO or 100 nM Tg for 30 minutes, then washed and replaced with normal media. Cellular RNA was harvested, 48 h post-treatment. Samples were sent for Illumina 3′ end bulk RNA-sequencing and gene set enrichment analysis (GSEA) was performed using the Reactome Pathway. Enrichment plot displaying up-regulation of genes related to cholesterol biosynthesis in Tg treatment vs. DMSO control. Normalized enrichment score (NES): 1.75, nominal p-value: <0.0001, FDR q-value: 0.016. **(B)** Huh7.5 cells were treated with DMSO or 100 nM Tg for 30 minutes and fixed at 4, 24, 48 and 72 h post-treatment. Cells were stained for cell nuclei (DAPI, blue) or LDs (BODIPY-493, green) and assessed by confocal microscopy under 63x magnification. **(C-D)** Huh7.5 cells were treated with DMSO or 100 nM Tg for 30 minutes followed by HCVcc infection (MOI 0.2) for 72 hours. **(C)** Cellular RNA was harvested and *FASN*, *DGAT1*, *DGAT2, ACAT1 and ACAT2* mRNA expression was measured by RT-qPCR. **(D)** Cells were fixed and stained for cell nuclei (DAPI, blue), LDs (BODIPY-493, green), or HCV core (red) and assessed by confocal microscopy under 63x magnification. **(E)** Average lipid droplet area was measured using the FIJI/image J image analysis software. Graphs show mean +/- SEM of at least three independent experiments performed in triplicate. Confocal microscopy data are representative of at least three biological replicates with at least three technical replicates per condition. *p<0.05, **p<0.01, ***p<0.0005, ****p<0.0001.

We assessed LD morphology following Tg treatment by performing a time course experiment where cells were treated with Tg (100 nM) or DMSO for 30 minutes, then washed and subsequently fixed at 4, 24, 48 and 72 h post-treatment. Fixed samples were stained using BODIPY-493, a neutral lipid stain, to visualize LDs. We observed that Tg increases *de novo* synthesis of LDs at early time points, and that enlarged LDs persist in Tg-treated cells relative to DMSO-treated cells for at least 72 h **(Figure 2B).**

We next validated the effect of Tg treatment on expression of genes related to LD biogenesis in HCV-infected cells at 72 hpi. We observed modest increases in *FASN* and *DGAT1* mRNA expression in HCV-infected cells that were pre-treated with Tg, but no difference in *SOAT1/hACAT1*, *DGAT2* and *SOAT2/ACAT2* **(Figure 2C)**. To examine the effect of Tg treatment on LDs during HCV infection, Huh7.5 cells were treated with DMSO or Tg (100 nM) for 30 minutes, and subsequently infected with HCVcc for 72 hours. Confocal microscopy revealed that cells pre-treated with Tg had a notable increase in enlarged LDs compared to cells treated with DMSO vehicle **(Figure 2D)**. Notably, there was detectable HCV core expression following Tg treatment, albeit in fewer cells compared to DMSO, which could reflect decreased spread of HCV. We further assessed the size of LDs in Tg-treated and control groups and found that the average area of LDs in Tg-treated cells is significantly greater than their DMSO counterpart **(Figure 2E)**.

### IRE1 pathway activation only partially recapitulates the antiviral effect of Tg against HCV

Activation of the IRE1 pathway of the UPR is associated with increased lipogenesis due to the transcriptional activity of Xbp1s. As such, we employed a selective activator of the IRE1 pathway, IXA4 (34), to determine if specific activation of IRE1 recapitulates the effect of Tg against HCV infection. In our cell model, treatment with IXA4 somewhat broadly activated all pathways of the UPR **(Figure 3A)**, with a minimal effect on cell viability **(Figure S2)**. IXA4 treatment modestly increased *FASN*, *DGAT1* and *DGAT2* mRNA expression **(Figure 3B)**, similar to Tg **(Figure 2C)**. We next tested the effect of IXA4 against HCV replication and infection. IXA4 treatment led to approximately a 70% reduction in HCV RNA replication using the HCV SGR model **(Figure 3C)**. During HCVcc infection, IXA4 treatment modestly increased the size and number of LDs, but did not affect the number of core-positive HCV-infected cells **(Figure 3D)**, in contrast to Tg **(Figure 2D)**. Consistently, IXA4 treatment had a relatively limited effect on HCVcc extracellular titer **(Figure 3E)** that likely reflects its modest inhibition of HCV RNA replication. Taken together, these data indicate that IRE1 pathway activation likely contributes to Tg-mediated inhibition of viral RNA replication, but it does not explain the full antiviral effect of Tg. Furthermore, treatment with IXA4 led to unexpectedly broad activation of all three pathways of the UPR, yet did not recapitulate the antiviral effect of Tg, suggesting the potential for UPR-independent antiviral mechanisms against HCV.

**Figure 3.**
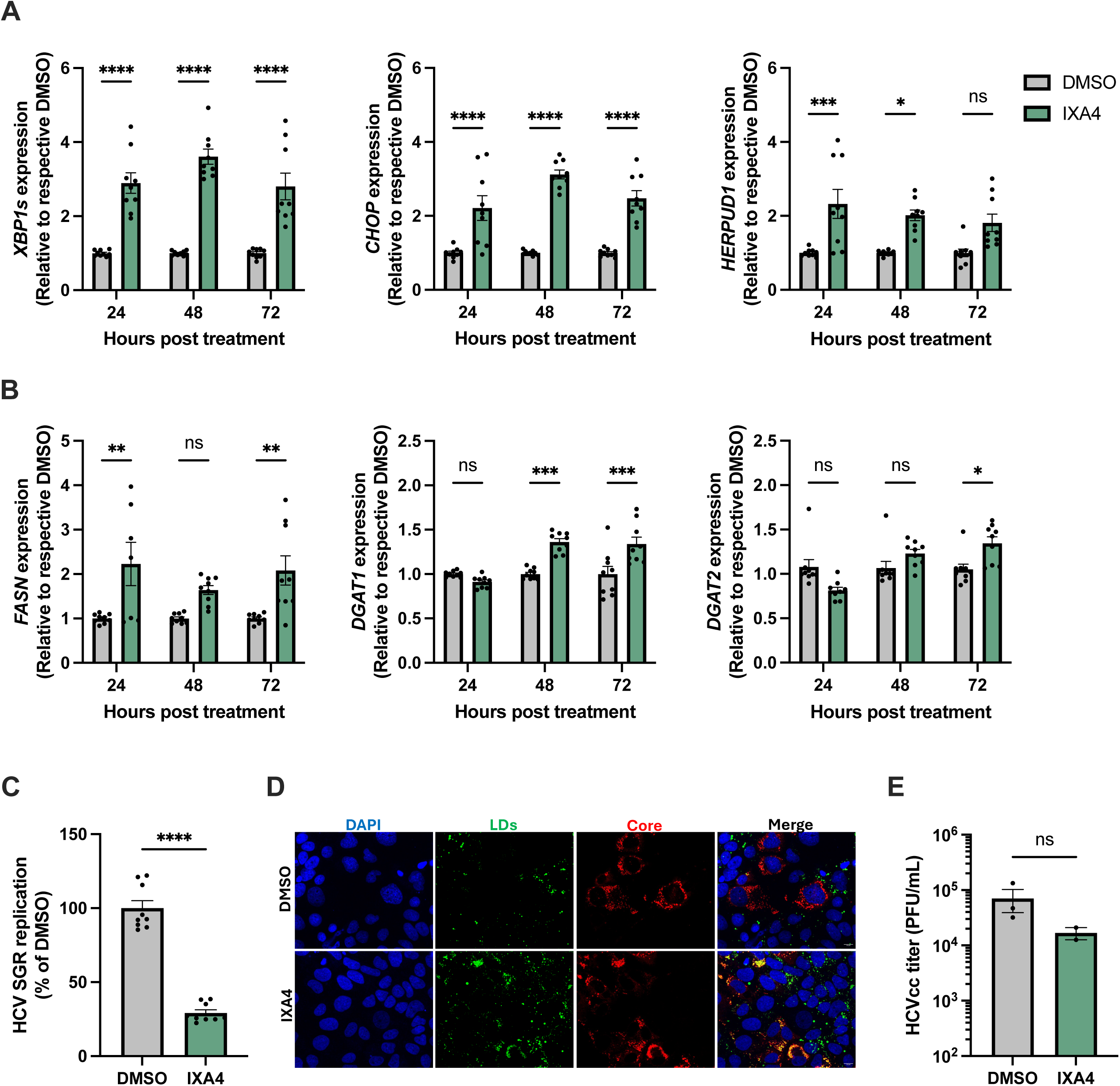
IXA4 treatment reduces HCV RNA replication and increases lipogenic gene expression without significantly affecting HCVcc titer. Huh7.5 cells were treated with DMSO or 10 µM IXA4 for 72 hours. **(A)** Cellular RNA was harvested and *XBP1s* (IRE1 pathway), *CHOP* (PERK pathway) and *HERPUD1* (ATF6 pathway) mRNA expression was assessed by RT qPCR. **(B)** Expression of genes related to LD biogenesis (*FASN*, *DGAT1* and *DGAT2*) was assessed by RT-qPCR. **(C)** Huh7 cells were electroporated with 5 µg of HCV SGR RNA. Four hours-post electroporation, cells were treated with DMSO or 10 µM IXA4 for 72h and RNA replication was measured by luciferase expression. Data are expressed as a percentage of HCV replication in DMSO-treated samples. **(D-E)** Huh7.5 cells were pre-treated with DMSO or 0.05 µM Tg for 30 minutes, and subsequently infected with HCV at MOI 0.2 for 72 h. **(D)** Cells were fixed and stained for cell nuclei (DAPI, blue) and LDs (BODIPY-493; green), HCV core (red), and assessed by confocal microscopy under 63x magnification. **(E)** Cellular supernatant was collected, and viral extracellular titer was assessed by focus forming assay. Graphs shown mean +/- SEM from three independent experiments. RT-qPCR was performed in technical triplicate per biological replicate. Confocal microscopy data are representative of at least three biological replicates with at least three technical replicates per condition. *p<0.05, **p<0.01, ***p<0.0005, ****p<0.0001.

### Tg-induced LD enlargement and inhibition of HCV infection is independent of the UPR

To further investigate the role of the UPR in the observed modulation of LDs and HCV infection, we used another UPR agonist, tunicamycin (Tm), that induces ER stress and the UPR by inhibiting N-linked protein glycosylation (35, 36). This results in the accumulation of misfolded proteins and induction of the UPR, but unlike Tg does not affect calcium flux. To confirm that pre-treatment with Tm induces ER stress and the UPR, we treated Huh7.5 cells with DMSO or Tm (1 µg/mL) for 4 hours and assessed UPR activation. As expected, treatment with Tm activated the UPR as measured by an increase in *XBP1s*, *HERPUD1* and *CHOP* expression **(Figure 3D)**. To assess the effect of Tm on LD morphology and HCV infection, we pre-treated cells with Tm for 4 hours prior to HCV infection (MOI 0.2) for 72 hours. Consistent with previous literature, Tm did not affect LD morphology, nor did it decrease the number of infected cells **(Figure 4B)**. Moreover, pre-treatment with Tm did not significantly inhibit HCVcc RNA replication **(Figure 4C)** or viral titer **(Figure 4D)**. Together, the results suggest that the antiviral activity of Tg against HCV is likely independent of the UPR.

**Figure 4.**
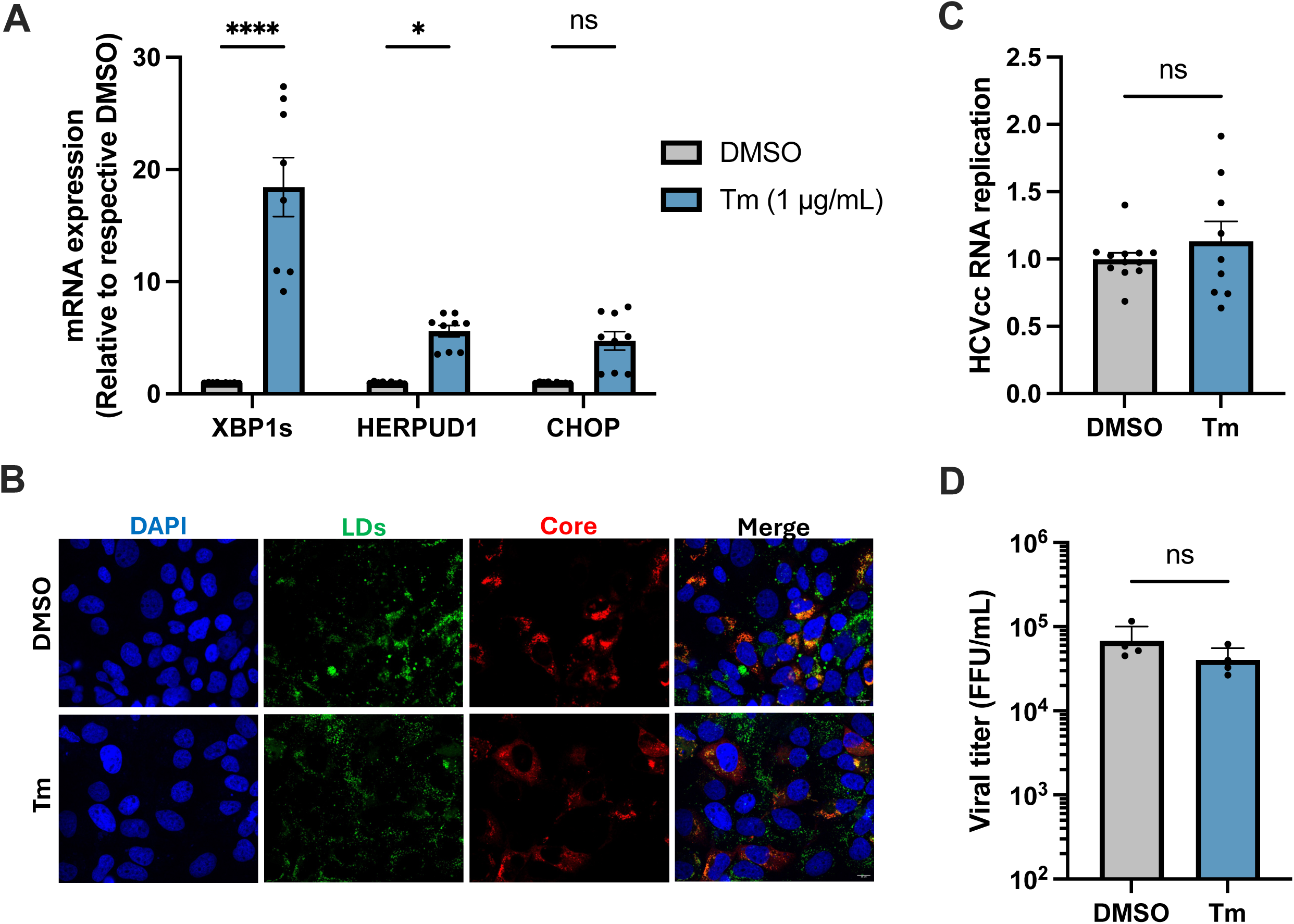
UPR activation by tunicamycin does not recapitulate the antiviral effect of Tg. (**A-C) (A)** Huh7.5 cells were treated with DMSO or 1 µg/mL tunicamycin (Tm) for four hours. Cellular RNA was harvested and *XBP1s* and *HERPUD1* mRNA expression was assessed by RT-qPCR. **(B-D)** Huh7.5 cells were pre-treated with DMSO or 1 µg/mL Tm for four hours, and subsequently infected with HCVcc (MOI 0.2) for 72 hours. **(B)** intracellular HCV RNA was assessed by RT-qPCR, **(C)** Cells were fixed and stained for cell nuclei (DAPI, blue) and LDs (BODIPY-493; green), HCV core (red), and assessed by confocal microscopy under 63x magnification. **(D)** Viral titer was assessed by focus forming assay. Graphs shown mean +/- SEM from three independent experiments. RT-qPCR was performed in technical triplicate per biological replicate. Confocal microscopy data are representative of at least three biological replicates with at least three technical replicates per condition. *p<0.05, **p<0.01, ***p<0.0005, ****p<0.0001.

### Tg-induced enlargement of LDs mediates inhibition of HCV RNA replication

Since we observed that Tg treatment leads to the accumulation of enlarged LDs **(Figure 2B,D)**, we hypothesized that Tg treatment may promote LD fusion. LD fusion is one mechanism through which LDs can mature and expand and is mediated by cellular proteins that localize to LD-LD contact sites. Cell death-inducing DFFA-like effector B (CIDE-B) and CIDE-C accumulate at LD-LD contact sites in hepatocytes to promote LD fusion (37). While CIDE-B is most abundantly expressed, CIDE-C is primarily associated with promoting large LD accumulation in hepatocytes (37). Localization of CIDE-C to LD-LD contact sites leads to the transfer of neutral lipids through leaky monolayers between LDs, resulting in LD fusion (38). Moreover, CIDE-C is transcriptionally regulated by metabolic transcription factors such as SREBPs, peroxisome proliferator-activated receptor gamma and CCAAT/enhancer binding protein (39) rather than canonical UPR sensor signalling. Thus, we next examined the role of CIDE-C in mediating the effects of Tg on LD morphology.

We first demonstrated that Tg treatment increases *CIDE-C* expression at the transcriptional level **(Figure 5A)**. Subsequently, we used RNAi to stably silence CIDE-C expression in Huh7.5 cells **(Figure 5B)** and assessed the effect on Tg-induced LD enlargement. Huh7.5 shCTRL or shCIDE-C cells were treated with DMSO or Tg (100 nM) for 30 minutes, and LD morphology was assessed 72 h post-treatment. Confocal microscopy analysis revealed that Tg-induced LD enlargement was significantly reduced in shCIDE-C cells compared to shCTRL cells **(Figure 5C)**. We next tested the effect of CIDE-C silencing on the antiviral activity of Tg against HCV. We first noted that HCV RNA replication was decreased by approximately 50% in Huh7.5-shCIDE-C cells compared to shCTRL cells **(Figure 5D)**. Interestingly, CIDE-C silencing appeared to rescue viral RNA replication from inhibition by Tg **(Figure 5D)**. However, despite rescuing viral replication, CIDE-C silencing did not fully rescue viral titer from the effects of Tg **(Figure 5E)**. These findings suggest that while Tg-mediated LD remodeling modestly impairs HCV RNA replication, additional antiviral mechanisms are responsible for its effects on viral titer.

**Figure 5.**
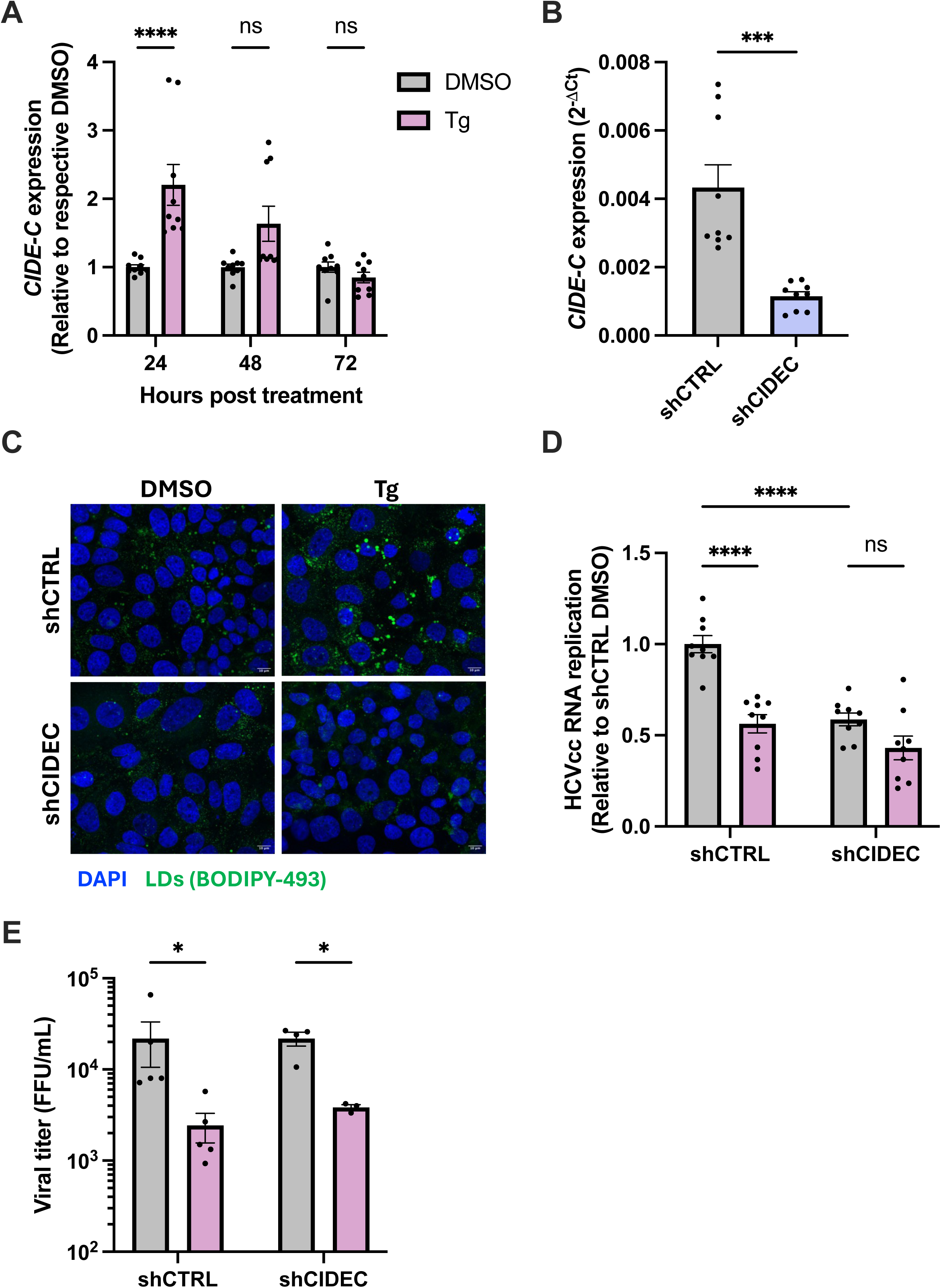
Silencing CIDE-C expression partially rescues Tg-induced HCV RNA replication. **(A)** Huh7.5 cells were treated with DMSO or Tg (100 nM) for 30 minutes, and cellular RNA was harvested 4, 24, 48 or 72 h post-treatment. *CIDE-C* mRNA expression was assessed by RT-qPCR. The data are normalized to actin and set relative to the DMSO control at each respective time point. **(B)** Huh7.5 cells were transduced with shCTRL or shCIDE-C lentivirus. Successful silencing of CIDE-C expression was confirmed by RT-qPCR. **(C)** Huh7.5 shCTRL or shCIDE-C cells were treated with DMSO or Tg (100 nM) for 30 minutes and fixed 72 hours post-treatment. Cells were stained for cell nuclei (DAPI, blue) or LDs (BODIPY-493, green) and assessed by immunofluorescence confocal microscopy under 63x magnification. **(D-E)** Huh7.5 shCTRL or shCIDE-C cells were pre-treated with DMSO or Tg (100 nM) for 30 minutes followed by HCVcc infection (MOI 0.2) for 72 h. Cellular RNA and supernatants were collected and **(D)** HCVcc RNA replication or **(E)** extracellular viral titer was assessed by RT-qPCR or focus-forming assay, respectively. Graphs shown mean +/- SEM from three independent experiments. RT-qPCR was performed in technical triplicate per biological replicate. Confocal microscopy data are representative of at least three biological replicates with at least three technical replicates per condition. *p<0.05, **p<0.01, ***p<0.0005, ****p<0.0001.

### Tg treatment disrupts HCV egress and ApoB and E secretion

Since silencing of CIDE-C expression did not rescue viral titer from Tg inhibition, we next sought to determine how Tg treatment affects later steps of the viral lifecycle. To discern if Tg affects virion assembly or secretion, we quantified intracellular viral titer. Huh7.5 cells were pre-treated with DMSO or Tg (100 nM) for 30 minutes and subsequently infected with HCV for 72h. Interestingly, we observed no difference in intracellular viral titer in Tg-treated cells, compared to the DMSO control. However, there was a significant decrease in extracellular titer **(Figure 6A)**, indicating that Tg does not affect viral assembly but likely affects secretion of progeny virions. To assess whether this reflected a defect in secretion or infectivity, we quantified extracellular viral RNA in DMSO or Tg-treated HCV-infected cells. Consistent with the decrease in extracellular titer, we observed a corresponding decrease in extracellular viral RNA **(Figure 6B)**, confirming that Tg treatment reduces the secretion of progeny virions.

**Figure 6.**
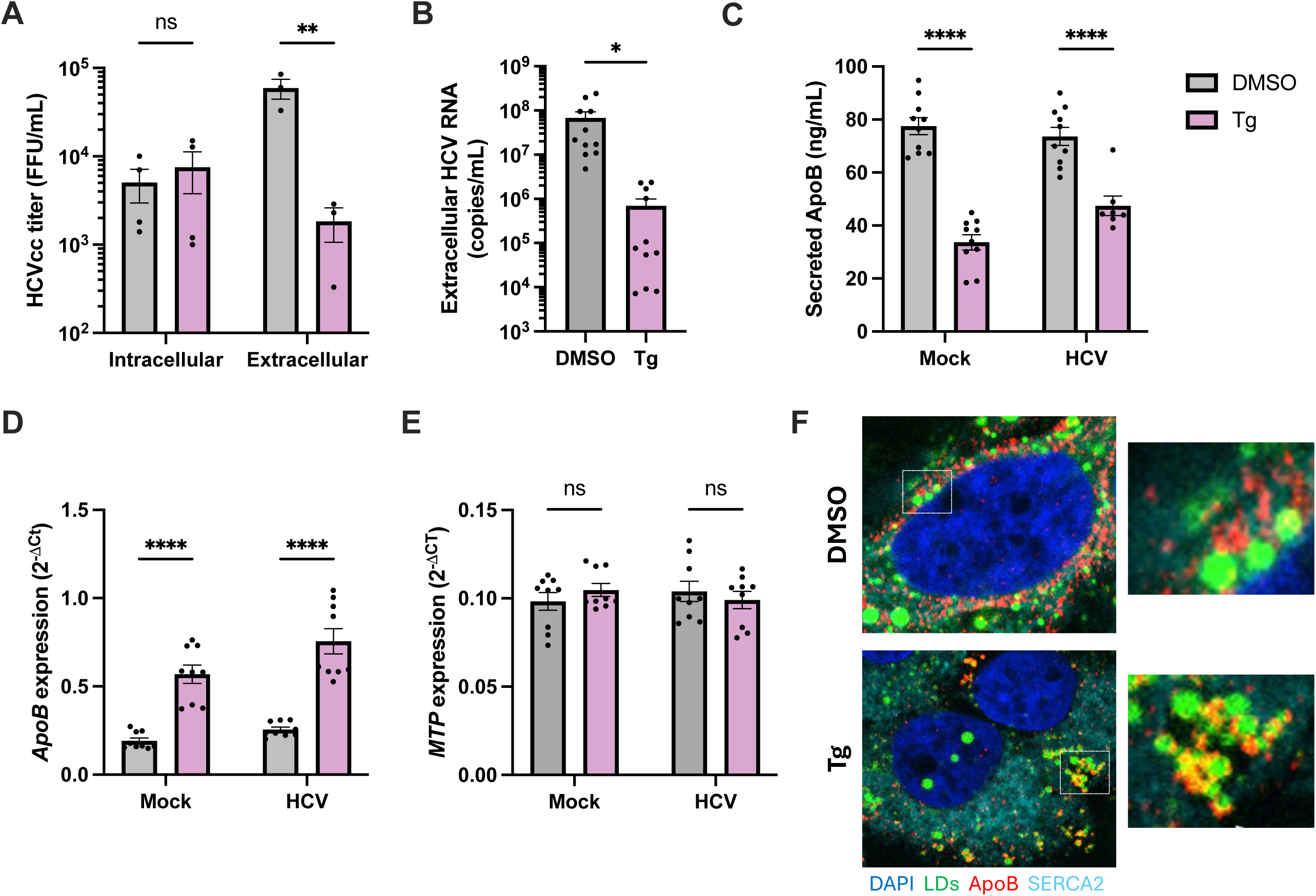
Tg treatment inhibits secretion of HCV and ApoB. **(A)** Huh7.5 cells were pre-treated with DMSO or 100 nM Tg for 30 minutes and subsequently infected with HCVcc (MOI 0.2) for 72 h. Cellular supernatants were collected to titrate extracellular virus, while cells were collected and subjected to 3 freeze-thaw cycles to recover intracellular virus. Viral titer was measured by focus-forming assay. **(B)** Infections were performed as described in **(A)**, and 72 hpi, extracellular viral RNA was isolated from supernatants. **(C-E)** Huh7.5 cells were pre-treated with DMSO or 100 nM Tg for 30 minutes and subsequently mock-infected or infected with HCVcc (MOI 0.2) for 72 h. **(C)** Cellular supernatants were collected and ApoB secretion was measured by ELISA or **(D)** ApoB and (E) MTP mRNA expression was assessed by RT-qPCR. **(F)** Mock infected Huh7.5 cells were treated DMSO or 100 nM Tg for 30 minutes, fixed 72 h post-treatment and stained for cell nuclei (DAPI; blue), SERCA2 (ER, cyan), ApoB (red) or LDs (green) and assessed by confocal microscopy under 60x magnification. Graphs shown mean +/- SEM from at least three independent experiments. ELISA was performed in technical duplicate per biological replicate. RT-qPCR experiments were performed in technical triplicate per biological replicate. Confocal microscopy data are representative of three biological replicates with at least three technical replicates per condition. *p<0.05, **p<0.01, ***p<0.0005, ****p<0.0001.

HCV classically egresses through the lipoprotein secretion pathway, where it interacts with various lipoproteins, including apolipoproteins (Apo) B and E, leading to the formation of lipoviral particles (40). It was previously reported that treatment with Tg decreased ApoB secretion in Huh7 cells (41). Indeed, ApoB secretion was significantly reduced in Tg-treated samples regardless of infection **(Figure 6C)**, suggesting that Tg may disrupt the very low-density lipoprotein (VLDL) secretion pathway that is required for efficient HCV egress. To validate that secretion, and not expression, of ApoB is affected by Tg treatment, we first assessed intracellular mRNA expression of ApoB by RT-qPCR. Huh7.5 cells were pre-treated with DMSO or Tg (100 nM) for 30 minutes and subsequently mock or HCV-infected for 72 h. Intracellular ApoB mRNA expression was upregulated in cells pre-treated with Tg compared to DMSO control **(Figure 6D**). ApoB co-translational lipidation is an essential step of VLDL biogenesis occurring at the ER and is mediated by the microsomal triglyceride transfer protein (MTP), in which triglycerides from LDs are transferred to ApoB within the ER lumen (42). Thus, we also assessed *MTP* mRNA expression but found that it was not affected by Tg treatment **(Figure 6E).** Finally, we evaluated ApoB protein expression and subcellular localization using confocal microscopy. If ApoB is properly synthesized and lipidated within the ER lumen but is unable to be exported from the ER to the Golgi, it has been described to form ApoB “crescents” along lipid droplets (43). To determine if ApoB is retained in the ER and forms these crescents, we stained for SERCA2 (ER marker), LDs and ApoB. ApoB expression appeared more diffuse in DMSO-treated cells, but we observed an increase in ApoB crescent formation on LDs following treatment with Tg **(Figure 6F).** Together, our data suggest that Tg may impair ApoB trafficking from the ER to the Golgi, leading to a defect in its secretion.

To more broadly assess the effect of Tg treatment on secretion, we next evaluated an array of lipoproteins, including ApoAI, ApoAII, ApoB-100, ApoCII, ApoCIII, ApoD, ApoE, ApoH, ApoJ and ApoM in mock or HCV infected cells using a cytometric bead array. Interestingly, we observed that Tg selectively impairs the secretion of ApoAI, ApoB-100 and ApoE in mock or HCV-infected cells **(Figure 7A).** While it is debated whether or not ER stress increases or decreases cellular secretory capacity (44), our bulk RNA-seq analysis showed that numerous genes associated with cellular secretion are upregulated following Tg treatment **(Figure 7B, Figure S1B)**. To functionally assess whether Tg treatment causes a specific defect in the secretion of lipoproteins or broadly disrupts cellular secretion, we utilized a Gaussia luciferase (Gluc) construct, which is rapidly secreted through the classical secretory pathway (45). Huh7.5 cells were transfected with pCMV-Gluc for 4 h and subsequently treated with DMSO or Tg for 30 minutes. We measured secreted and intracellular Gluc expression at 24 h following Tg treatment and calculated the secretion ratio of extracellular Gluc signal to intracellular Gluc signal. As expected, brefeldin A (BFA), which is well-established to disrupt the secretory pathway (46), decreased the Gluc secretion ratio **(Figure 7C).** Interestingly, the Gluc secretion ratio was not decreased, but rather increased in cells treated with Tg compared to DMSO **(Figure 7C)**, reflecting the upregulation of protein secretion that we observed in our RNA-Seq analysis **(Figure 7B)**. Taken together, our data show that Tg treatment does not broadly disrupt cellular secretory function, but selectively disrupts the secretion of HCV, likely through specific modulation of the VLDL secretion pathway.

**Figure 7.**
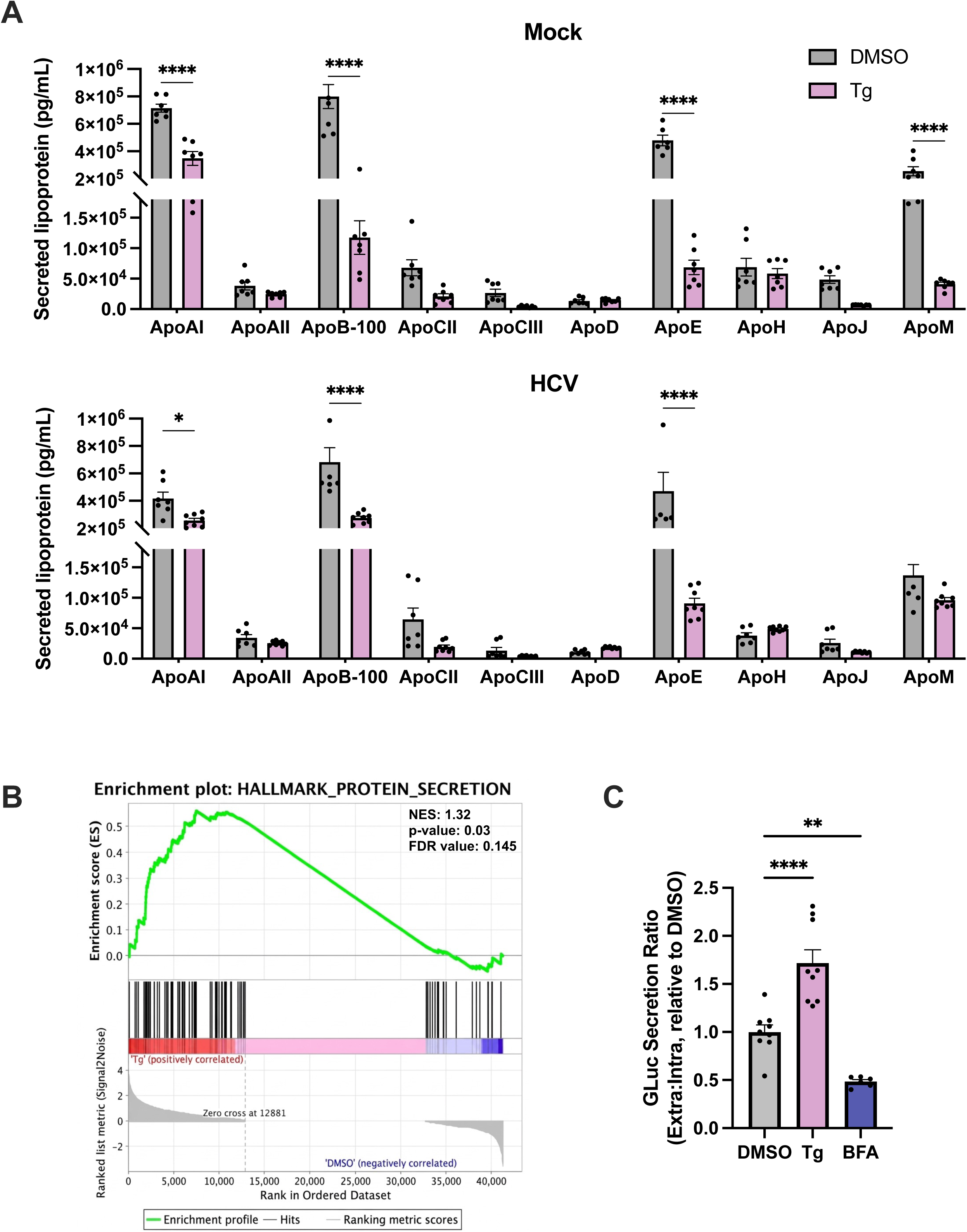
Treatment with Tg selectively disrupts the secretion of certain lipoproteins. (**A**) Huh7.5 cells were pre-treated with DMSO or 100 nM Tg followed by mock-infection or infected with HCVcc (MOI 0.2) for 72 h. Supernatants were harvested and the expression of various lipoproteins (ApoAI, ApoAII, ApoB-100, ApoCII, ApoCIII, ApoD, ApoE, ApoH, ApoJ and ApoM) was assessed using a cytometric bead array. (B) Huh7.5 cells were treated with DMSO or 100 nM Tg for 30 minutes, then washed and replaced with normal media. Cellular RNA was harvested, 48 h post-treatment. Samples were sent for Illumina 3′ end bulk RNA-sequencing and gene set enrichment analysis (GSEA) was performed using the Hallmark Pathway. Enrichment plot displaying up-regulation of genes involved in protein synthesis in Tg treatment vs. DMSO control. Normalized enrichment score (NES): 1.32, nominal p-value: 0.03, FDR q-value: 0145. (C) Huh7.5 cells were transfected with 500 ng pCMV-Gluc for 4 h and subsequently treated with DMSO or 100 nM Tg for 30 minutes. Intracellular and secreted Gluc expression was measured 24 h post treatment. Data are shown as a secretion ratio of extracellular Gluc: intracellular Gluc. Graphs shown mean +/- SEM from at least three independent experiments. Multiplex assay was performed in technical duplicate per biological replicate. Luciferase experiments were performed in technical triplicate per biological replicate.

## DISCUSSION

While the antiviral mechanism of Tg has remained elusive, our recent study has suggested that its antiviral activity against human coronavirus infection may be independent of the UPR (13). Here, we aimed to investigate potential alternative antiviral mechanisms, namely the role of Tg-induced perturbations to lipid metabolism, using HCV as a model. We report that a short pre-treatment of cells with Tg is sufficient to inhibit HCV infection. We characterized the effect of Tg at various steps of the HCV replication cycle and show that the greatest effect is on viral titer, indicating a block to late stages of infection, such as assembly or secretion.

Given the key role of LDs in mediating HCV nucleocapsid assembly (22–24), we focused on how Tg alters LD morphology and lipid homeostasis and the impact this has on HCV infection, which could explain the decreased viral titer observed in Tg-treated cells. Consistent with previous literature (18), Tg treatment upregulated the expression of genes involved in LD biogenesis, such as *FASN, DGAT1* and *DGAT2*. In parallel, we observed a higher frequency of enlarged LDs in Tg-treated samples, with enlarged LDs persisting for at least 72 hours post treatment. Similarly, Leal et al. reported that HepG2 cells treated with various concentrations of Tg (50-200 nM) for 24 hours was sufficient to cause an increase in lipid accumulation (47), which was attributed to UPR activation.

While all three pathways of the UPR contribute to maintain lipid homeostasis, the IRE1 pathway is most well described to promote lipogenesis. Activation of IRE1 results in its dimerization and autophosphorylation, leading to the activation of the IRE1 endoribonuclease domain. The endonuclease activity of IRE1 mediates the non-conventional cytoplasmic splicing of X-binding protein 1 (*XBP1*), removing a 26-nucleotide intron, resulting in a spliced *XBP1* variant, *XBP1s* (12). XBP1s is a potent transcription factor that regulates the expression of key lipogenic genes, including fatty acid synthase (*FASN*) and diacyl glycerol acetyltransferase 2 (*DGAT2*) (48). Moreover, knockout of XBP1s in the murine liver resulted in a decreased production of lipids (48), highlighting the importance of this pathway in regulating lipid metabolism. Therefore, we evaluated the effect of IRE1 activation on HCV infection. Treatment with IXA4, which activates the endoribonuclease domain of IRE1, caused a significant reduction in HCV SGR replication. However, in the context of HCVcc infection, IXA4 treatment had a minimal effect on core expression, viral spread, and viral titer. IXA4 treatment resulted in a modest increase in lipogenic gene expression and LDs but did not recapitulate the effect of Tg. Thus, activation of the IRE1 pathway may contribute to the observed inhibition of RNA replication mediated by Tg. Tardif et al. reported that cells stably expressing the HCV SGR exhibit decreased XBP1s transcriptional activity and ER-associated degradation, two canonical downstream effects of IRE1 activation (49), suggesting that HCV infection suppresses the IRE1-XBP1s pathway. Thus, pre-activation of the IRE1 pathway through IXA4 or Tg treatment may overcome the ability of HCV to antagonize these responses, leading to the antiviral effect we observed. It is worth noting that while IXA4 has previously been described as a selective activator (34), we observed relatively broad UPR activation, reflecting the overlap and crosstalk between the three UPR pathways. Future studies in which XBP1s or a pre-activated form of IRE1 is overexpressed would be helpful to better understand the specific contribution of the IRE1 pathway independently of other UPR pathways.

While we observed that IXA4-induced ER stress partially inhibits HCV SGR replication, this effect was not reflected in HCVcc infection, indicating that UPR activation alone may not be responsible for the full antiviral effect of Tg. Indeed, our recent study assessing the antiviral activity of Tg against a human common cold coronavirus 229E (HCoV-229E), revealed that expression of any of the UPR sensors is not required for Tg-mediated inhibition of HCoV-229E replication (13). Consistently, Tm, which induces ER stress and the UPR by disrupting N-linked glycosylation, did not inhibit HCV infection. We observed no differences in intracellular RNA expression in HCV-infected cells, which is in line with previous results showing that treatment with Tm did not reduce NS5A expression in cells expressing the HCV SGR (50). Furthermore, Tm treatment did not overtly affect LD morphology in HCV-infected cells. This is consistent with previous reports that Tg, but not Tm, modulates LDs during SARS-CoV-2 pseudoparticle entry (41).

Given the importance of glycosylation of HCV E1 and E2 glycoproteins during egress, the limited effect of Tm on viral titer is surprising. It is possible that following the removal of Tm, treated cells recover more rapidly and normal glycosylation is at least partially resumed prior to egress of HCV particles. In contrast, Tg priming causes a prolonged effect on the cell, reflecting its irreversible inhibition of SERCA (51) and the long protein half-life of SERCA (52, 53). Overall, UPR activation by Tm is not sufficient to induce the observed modulation of LDs following Tg priming, pointing to a potential UPR-independent mechanism for Tg. For example, Tg treatment could alter LD morphology through depletion of calcium by inhibition of SERCA (51) or through modulation of ER-membrane cholesterol content, which can activate SREBP1/2 (54, 55), leading to an increase in lipogenic gene expression that may explain the observed effect on LDs and the antiviral phenotype.

We next sought to further understand whether enlargement of LDs in Tg-primed cells restricts HCV infection. Silencing the expression of CIDE-C, a mediator of LD fusion, reduced Tg-induced LD enlargement, and rescued viral RNA replication, but not viral titer, from inhibition by Tg. Modulation of lipid metabolism is well-documented to inhibit HCV infection. Treatment with lavostatin (HMG-CoA reductase inhibitor) or 2-chloro-5-nitro-N-(pyridyl)benzamide (BA) (FASN inhibitor) is reported to inhibit HCV replication in Huh7.5 or Huh7 replicon expressing cells (56–58). Thus, Tg-induced alterations in lipid metabolism could similarly impact viral RNA replication. HCV ROs are highly enriched in cholesterol (59). One possibility is that induction of enlarged LDs by Tg results in more cholesterol being sequestered in LDs, thereby decreasing available cholesterol for ROs. Blocking Tg-induced LD enlargement through silencing CIDE-C expression may promote a greater localization of cholesterol and other lipids to sites of viral RNA replication, although this remains to be experimentally tested. Notably, another study found that overexpression of DGAT2 increased the size of LDs during HCV infection and strongly suppressed RO formation and HCV replication (60). DGAT2 overexpression and HCV infection increased the levels of similar types of lipids that promote membrane fluidity and curvature, which are required for both RO and LD formation, resulting in a competition for lipids (60). As such, it is possible that Tg treatment similarly induces a competition for lipids between ROs and LDs, which negatively impacts HCV RO formation and thus viral RNA replication. It is worth noting that silencing CIDE-C expression reduced HCV RNA replication, which may be due to increased LD fragmentation (via lipolysis) and disruptions in triglyceride homeostasis (61) that could negatively affect HCV infection. It is also possible that depletion of CIDE-C may alter the lipid composition or proteome of LDs to promote an enhanced antiviral state. Nevertheless, CIDE-C silencing did not affect the ability of Tg to decrease viral titer.

Our further characterization of the inhibitory effect of Tg on later stages of the viral lifecycle revealed that Tg does not significantly intracellular HCV titer, despite a significant decrease in both extracellular titer and RNA, suggesting a block to the secretion. Secretion of HCV particles is closely linked to the VLDL secretion pathway, which is largely dependent on ApoB (62). In Tg-treated mock or HCV-infected cells, we observed a significant decrease in ApoB secretion, suggesting a potential blockade to the host-lipoprotein secretion pathway. This is in line with a previous study demonstrating that Tg treatment decreases ApoB secretion during SARS-CoV-2 pseudoparticle entry (41). Indeed, inhibition of MTP and ApoB lipidation in Huh7 cells containing an integrated genotype 2a cDNA (Huh7-GL), which allow the production of infectious virus, resulted in a significant decrease in extracellular viral titer and extracellular viral RNA, but had no effect on intracellular RNA expression (62). Moreover, silencing of ApoB had minimal effect on intracellular viral RNA expression, but reduced extracellular titer (62). These findings reflect similar trends observed in our study. Consistently, silencing of ApoB expression resulted in a ∼70% decrease in ApoB secretion, a 75% decrease in core secretion and a ∼70% decrease in positive strand HCV RNA secretion (63).

Inhibition of MTP by the flavonoid naringenin decreased secretion of HCV particles by ∼80% (63). While we observed no decrease in the mRNA expression of MTP, it is possible that Tg treatment decreases MTP protein expression or enzymatic activity which would impair ApoB lipidation (promoting ApoB degradation) and VLDL assembly, explaining the defect we observed on both ApoB and HCV secretion. In contrast, a previous study showed that a 4 day treatment with Tg (100 nM) significantly decreases *MTP* mRNA expression in HepaRG cells (64). The lack of effect on MTP mRNA expression in our studies could be explained by the difference in the duration of Tg treatment or cell model. Following Tg treatment, there is an increase in ApoB mRNA expression with no significant difference in protein expression. However, we observed changes in the subcellular localization of ApoB, where ApoB is more localized to LDs in Tg-treated cells, reflected by an increase in the formation of ApoB crescents along LDs. This observation suggests that there is a block to the anterograde transport of ApoB from the ER to the Golgi, thereby inhibiting ApoB secretion (43).

To assess the specificity of Tg in blocking secretion, we evaluated the effect on the secretion of an array of lipoproteins, and found that in addition to ApoB, secretion of ApoAI and ApoE is disrupted, while secretion of all other tested lipoproteins was not significantly affected. Notably, Tg caused a strong decrease in the secretion of ApoB-100 and ApoE, which are the most relevant lipoproteins for HCV. ApoE associates with HCV lipoviral particles and is required for virion morphogenesis and infectivity by mediating binding of virions to heparan sulfate proteoglycans, low density lipoprotein receptor, very low density lipoprotein receptor and scavenger receptor B1 (65). Interestingly, Hishiki and colleagues reported that when ApoE secretion is disrupted by expressing an ApoE construct that is retained in the ER, the secretion of infectious HCVcc is decreased and is instead retained intracellularly (66). Our data phenocopies this effect, wherein treatment with Tg disrupts ApoE secretion without affecting intracellular accumulation of infectious viral particles. It is worth noting that one study has reported that ApoA1 can in part rescue HCV infectivity in cells where ApoB and ApoE are knocked out (67). Ultimately, we propose that Tg-mediated inhibition of ApoB and ApoE secretion impairs the release of HCVcc.

VLDLs and low-density lipoproteins are large, bulky cargo that cannot fit in standard coat protein complex II (COP-II) vesicles that mediate anterograde trafficking and thus require specialized machinery to be transported from the ER to the Golgi (68). While the exact mechanism through which Tg disrupts ApoB and ApoE secretion requires further investigation, is possible that Tg treatment decreases ER exit sites, disrupts successful COP-II vesicle formation (69), or decreases the expression of adaptor proteins that are specifically required for the transport of larger cargo, such as TANGO1 (68). Moreover, it was reported that treatment with Tg (Huh7 cells, 300 nM) decreases the anterograde trafficking of temperature sensitive vesicular stomatitis virus glycoprotein (VSV-G) (69). To complement these experiments using a non-lipid dependent protein, we leveraged a Gluc expression construct which is rapidly secreted through the canonical secretory pathway. Treatment with Tg enhanced Gluc secretion compared to DMSO-treated cells, in line with a previous study showing that induction of ER stress can increase the secretion of certain proteins as a mechanism to restore extracellular proteostasis (70). Thus, a short treatment with Tg may enhance constitutive secretory pathway function or alter cellular trafficking, as our data suggests. Overall, the increase in Gluc secretion coupled with the decrease in ApoB and ApoE secretion suggests that Tg treatment selectively impairs VLDL secretion. Ultimately, we propose that Tg-mediated inhibition of ApoB and ApoE secretion impairs the release of infectious HCVcc particles.

In conclusion, our data demonstrates that Tg inhibits HCV infection through distinct mechanisms resulting from alterations to lipid metabolism that inhibit both HCV RNA replication and assembly. Tg-induced LD enlargement modestly inhibits HCV RNA replication, but more profoundly inhibits HCV egress by impairing VLDL secretion. The modulation of lipid metabolism by Tg has implications for many RNA viruses. For example, SARS-CoV-2 replication organelles are closely associated with LDs (71, 72), and thus disruption to LDs may restrict RNA replication and could contribute to the observed inhibition of CoV RNA replication by Tg (13). Moreover, since DENV and ZIKV rely on LDs to fuel virion morphogenesis (55, 73), it is possible that Tg-induced alterations in the size or composition of LDs could alter virion formation and maturation, explaining the previously reported inhibition of these viruses by Tg. Overall, modulation of lipid metabolism and selective impairment of secretory trafficking by Tg represents a novel antiviral mechanism which could have broader implications for other RNA viruses that rely on host lipid metabolism to support infection.

## MATERIALS AND METHODS

### Mammalian cell lines

Huh7 cells (JCRB0403) were obtained from the Japanese Collection of Research Bioresources Cell Bank. Huh7.5 (APC49) and Huh7.5.1 (APC167) cells were obtained from Apath LLC. 293T/17 (CRL-11268) cells were obtained from American Type Culture Collection. Huh7, Huh7.5, Huh7.5.1 and 293T/17 cells were cultured in Dulbecco’s Modified Eagle Medium (DMEM; Thermo Scientific #11995065) supplemented with 10% FBS (Corning, #35-077-CV) and 1% penicillin/streptomycin (Thermo Scientific, #15140122).

### HCV propagation and virus infection

The plasmid encoding the full length HCV Japanese Fulminant Hepatitis-1 (JFH-1_T_) (74) was kindly provided by Dr. Rodney Russell (Memorial University). The JFH1_T_ plasmid was linearized through restriction digestion with XbaI and subjected to a DNA clean up using the Monarch PCR and DNA cleanup Kit (#T1030S) as per the manufacturer’s protocol. The linearized plasmid DNA was in vitro transcribed using the NEB HiScribe High Yield T7 in vitro transcription kit (New England Biolabs, #E2040S) as outlined in the manufacturer’s protocol. The in vitro transcribed RNA was recovered using the Monarch RNA cleanup kit (#T2050S), aliquoted and stored at –80°C for subsequent use.

The HCV RNA was electroporated into 2 x 10^6^ Huh7.5 cells using the Neon electroporation system (Thermo Scientific). Following electroporation, the cells were plated into a 10-cm round dish and media was replaced with fresh DMEM containing 10% FBS 4 hours later. At 72 hours post-electroporation, supernatant was collected, spun at 500 x g for 4 minutes to remove cellular debris and filtered through a 0.45 µm filter. Virus was aliquoted, stored at –80°C and subsequently used for large scale HCVcc propagation.

For large scale HCVcc propagation, Huh7.5 or Huh7.5.1 cells were seeded in a 6-well plate at 3.0 x 10^5^ cells/well. The next day, cells were infected with HCVcc (MOI ∼0.003) diluted in serum-free media. After 4 h, inoculum was removed, and media was replaced with DMEM containing 10% FBS (Day 0). Cells were split and expanded on days 3 and 6, and media containing virus was collected on day 8. Supernatant was centrifuged at 500 x g for 5 minutes to remove cellular debris and subsequently filtered through a 0.45 µm filter. To concentrate virus, filtered supernatants were centrifuged through a Pierce Protein Concentrator with a 100K molecular weight cut off (Thermo Scientific, #88532) at 3000 x g for 15 minutes. The remaining virus was collected, aliquoted and stored at –80°C.

All virus infections were performed at an MOI of 0.2 focus-forming units (FFU) per cell. Briefly, cells were pre-treated with DMSO or Tg (100 nM) for 30 minutes and subsequently washed with 1X PBS. Viral inoculum was diluted in serum-free DMEM, and cells were infected for 4 h with occasional rocking of the plate. After 4 h, inoculum was removed and replaced with DMEM containing 2% FBS. Unless otherwise stated, infections were completed after 72 h.

### Virus titrations by focus forming assay

Cell culture supernatants were collected from infected cells and stored at –80°C. For intracellular titer, infected cells were pelleted, resuspended in DMEM (2% FBS, 1% P/S), and subjected to three freeze-thaw cycles from –80°C to 37°C. Viral titer was assessed by focus forming assay. Briefly, Huh7.5 or Huh7.5.1 cells were seeded at a density of 1 x 10^5^ cells/well in a 24-well plate. The next day, cellular supernatants containing HCV were serially diluted in serum-free DMEM. Cells were washed with serum-free DMEM and infected with 150 µL diluted virus at 37°C for four hours. Viral inocula were removed and replaced with DMEM containing 10% FBS and 1% P/S and incubated for 72 hours. Infected cells were fixed with 10% formalin for 1 h. Fixed cells were blocked and permeabilized in PBS containing 5% FBS and 0.1% Triton-X100 (blocking buffer) for one hour. Cells were stained with primary antibody against HCV core for 1 hour and subsequently stained with secondary antibody for 45 minutes. Foci of infected cells were counted using a Nikon Eclipse Ts2-FL inverted microscope.

### Drug treatments

Thapsigargin was purchased from Thermo Scientific (#T7458). Tunicamycin was purchased from Sigma Aldrich (T7765). IXA4 (#S9797) was purchased from Selleckchem. All stocks were prepared in dimethyl sulfoxide (DMSO; Thermo Scientific, # D12345), which was utilized as the vehicle control.

### Plasmids

All plasmids were propagated in NEB Stable Competent *Escherichia coli* (#C3040H). The pCMV-Gaussia luciferase construct was generously provided by Dr. Craig McCormick (Dalhousie University, Canada). The pLKO.1 plasmid (#26655) was purchased from Addgene. The pLKO.1 control plasmid was generated using the following non-targeting (scramble) sequence: 5’-CCTAAGGTTAAGTCGCCCTCG-3’. pLKO.1-shCIDEC was generated using the following sequence: CIDEC: 5’-AACTGTAGAGACAGAAGAGTA-3’. For production of lentivirus, pCMV-VSV-G (#8454) and psPAX2 (#12260) were purchased from Addgene. For production of HCV pseudoparticles, psPAX2 was used along with a pLenti-CMV-puro-Luc luciferase-encoding lentiviral genome (Addgene, #17477), and the pD603-H77-E1E2 plasmid encoding the H77 E1 and E2 glycoproteins (Addgene, #86983). The plasmid encoding the HCV subgenomic replicon (30) was kindly provided by Dr. Ralf Bartenschlager (Heidelberg University).

### Lentivirus production

Lentivirus for stable knockdown of shCTRL and shCIDEC in Huh7.5 cells was produced by seeding 293T/17 cells at a density of 8 x 10^5^ cells/well in a 6-well plate overnight. Cells were subsequently transfected with the following mixes: 600 ng psPAX2 packaging plasmid, 800 ng pLKO.1 transfer plasmid, 500 ng pCMV-VSV-G envelope plasmid, and 6 µL of lipofectamine 2000 (Thermo Scientific, #11668027). Supernatants were collected at 48 and 72 h post-transfection and filtered with a 0.45 µm filter. Lentivirus was aliquoted and stored at –80°C until further use.

For the production of HCVpp, HEK 293T/17 cells were seeded in 6-well plates at 5 x 10^5^ cells/mL and incubated overnight. The next day, cells were transfected with the following transfection mixture: 300 ng of psPAX2 (packaging plasmid), 500 ng of pLentiLuc, 300 ng of pD603-H77-E1E2, and 3 µL Fugene 6 transfection reagent up to a total of 100 µL in OptiMEM. The mixture was incubated at room temperature for 15-20 minutes and subsequently added dropwise to each well containing complete DMEM. After 16 hours, media was replaced with fresh complete DMEM. Supernatants containing the HCVpp were collected 48 and 72 h post-transfection, filtered through a 0.45 µm filter, aliquoted and stored at –80°C.

### Generation of stable cell lines

To generate stable cell lines by lentivirus transduction, Huh7.5 cells were seeded in a 6-well plate at 2 x 10^5^ cells/well overnight. The next day, cells were transduced with lentivirus in the presence of 8 µg/mL polybrene. Transduced cells were selected 72 h post-transduction using either 10 µg/mL blasticidin or 3 µg/mL puromycin. Successful knockdown was confirmed by RT-qPCR.

### Antibodies

For immunofluorescence confocal microscopy, cells were stained for HCV core (Anogen, #MO-I40015B, dilution 1:200), SERCA2 (Thermo Scientific #MA3-919, dilution 1:100), apolipoprotein B (Proteintech, #20578-1-AP, dilution 1:50) or with a neutral lipid dye, BODIPY-493/503 (Thermo Scientific #D3922, 2 µg/mL).

### RNA Isolation and RT-qPCR

Cellular RNA was harvested by lysing cells with 350 µL of RNA lysis buffer from the Monarch Total RNA Miniprep Kit (New England Biolabs, #T2110) or the QIAGEN RNeasy kit (QIAGEN, #74106) as per the manufacturer’s protocol. Cellular supernatants containing HCV were collected and stored at –80°C. Extracellular viral RNA was isolated using the Invitrogen Purelink viral RNA/DNA mini kit (Thermo Scientific, #12280050) as per the manufacturer protocol. Isolated RNA was reverse transcribed to generate cDNA using the High-Capacity cDNA synthesis kit (Thermo Scientific, #4368814). Quantitative PCR (qPCR) was performed using the PowerTrack SYBR Green Master Mix (Thermo Scientific, #A46111) using specific primers outlined in **Table 1**. Cellular gene expression and intracellular viral RNA levels were normalized to actin using the 2^-ΔCt^ method. For extracellular viral RNA, the number of genome copies was determined by comparing to a standard curve generated using an JFH-1_t_-encoding plasmid of known genome copy number.

**Table 1.**
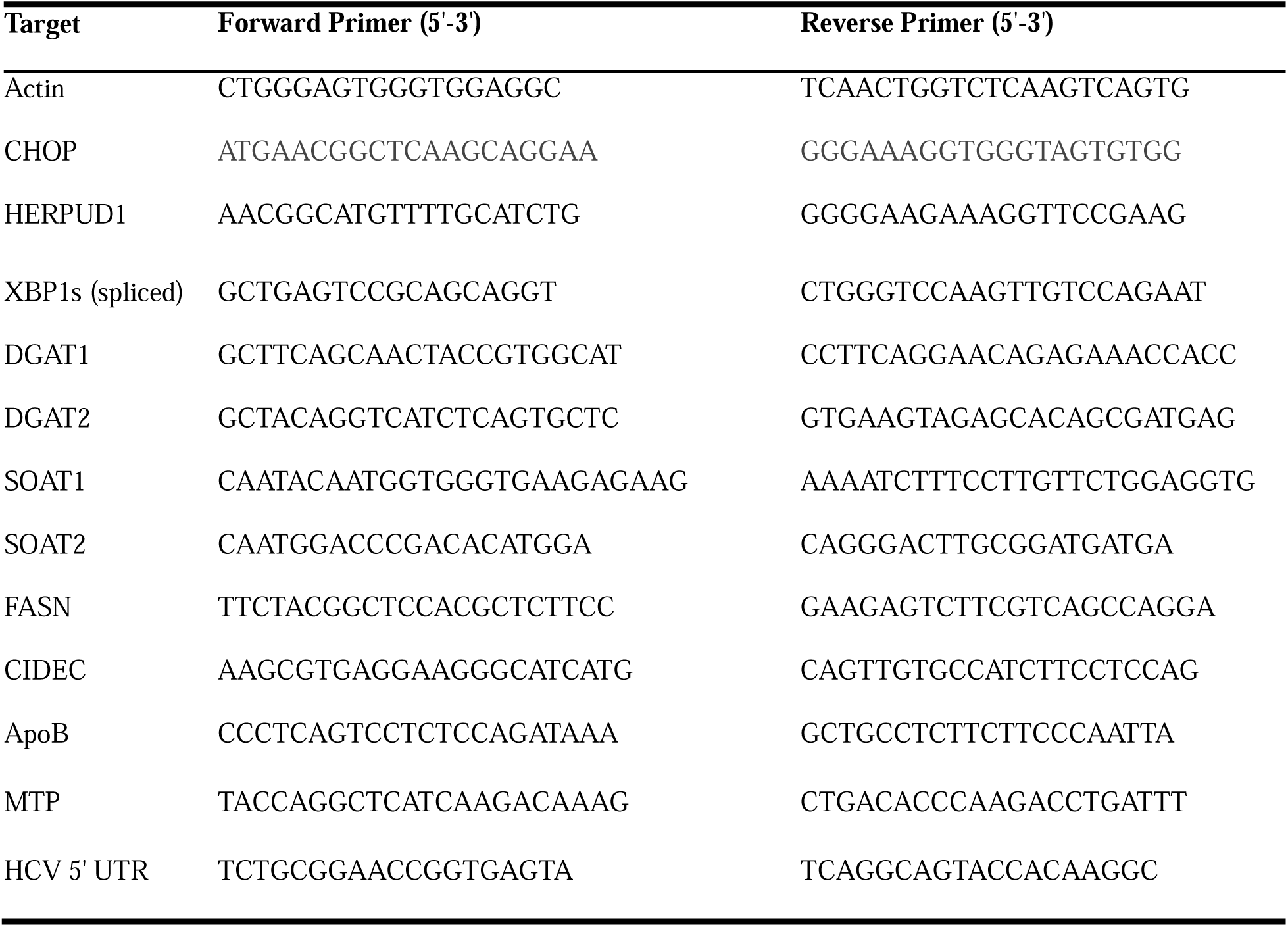
Oligos used in this study.

### Bulk RNA sequencing and analysis

Cellular RNA was harvested by lysing cells with 350 µL of RNA lysis buffer from the QIAGEN RNeasy kit (QIAGEN, #74106) supplemented with 1% Beta-mercaptoethanol. RNA was isolated/purified as per the manufacturer’s protocol with the optional DNAseI treatment (QIAGEN, #79254). Samples were submitted to Plasmidsaurus (Louisville, KY, USA) and prepared for sequencing according to the Plasmidsaurus standard workflow. Sequencing was performed using Illumina 3’-end RNA sequencing, generating deduplicated 3’-end gene reads for downstream analysis. The resulting gene expression profile table was uploaded to the desktop gene set enrichment analysis software (75) or the data were evaluated using pathway analysis provided by Plasmidsaurus. Genes of interest were selected based on pathway enrichment analysis. The counts per million (CPM) for each gene of interest was transformed into Log2 (CPM + 1) and their corresponding Z score was calculated. Z score values were uploaded to the Broad Institute Morpheus software for generation of a heatmap.

### Immunofluorescence

Cells were seeded on glass coverslips at 2 x 10^5^ cells/well in a 12-well plate. The next day, cells were treated with 100 nM Tg for 30 minutes, and either mock-infected or infected with HCV at an MOI of 0.2 ffu/cell. Cells were fixed in 10% formalin solution for 1 hour at room temperature at the corresponding time points. Fixed samples were blocked in PBS containing 5% FBS, 1% BSA and 0.01% saponin for one hour. Cells were stained for HCV core in PBS/1%BSA/0.01% saponin solution for two hours, followed by staining with AlexaFluor-555 or AlexaFluor-594 secondary antibody. Lipid droplets were stained using 2 µg/mL BODIPY-493 diluted in PBS for 30 minutes at room temperature, protected from light. Glass coverslips were mounted on microscope slides using FluoroMount G with DAPI (Thermo Scientific, #501128966) and visualized under 63x magnification under water immersion using a Leica Mica confocal microscope or under 60x under oil immersion using the Nikon-AX confocal microscope.

### Lipid droplet area measurement

Lipid droplet area was measured using FIJI image analysis software. Acquired images were set to 8 bit, and the background was subsequently subtracted. The threshold for each image was set individually due to significant changes in LD size across samples. To separate large LDs, the watershed feature was employed. LD particle size was subsequently analyzed using the following parameters: particle size: 0.2-infinity, circularity: 0,2-1. The generated table was copied into excel and the average area measurement was taken.

### In vitro transcription and electroporation of HCV subgenomic replicon RNA

Plasmid DNA was linearized by digestion with MluI and in vitro transcribed using the T7 MEGAscript Kit as per the manufacturer protocol. RNA was resuspended in nuclease-free dH_2_O at a concentration of 1 µg/µL and stored at –80°C. HCV SGR RNA was electroporated into 2 x 10^6^ Huh7 cells using the Neon transfection system (Thermo Scientific). Huh7 cells were pelleted, resuspended in 100 µL of Buffer R, and combined with 5 µg of HCV SGR RNA, then loaded into a Neon tip and electroporated using the following parameters: 1400 V, 20 ms, one pulse. Electroporated cells were resuspended in pre-warmed DMEM-10% FBS and subsequently seeded into 96-well plates at a density of 2.0 x 10^4^ cells/well. At 4 h post-electroporation, cells were treated with DMSO or 100 nM Tg (30 minutes) or 10 µM IXA4. Luciferase expression was assessed at 24, 48 and 72 h post-electroporation. Cells were lysed using 0.1% Triton-X100 and combined with firefly luciferase reagent (Nanolight #318) and transferred to a white flat-bottom 96-well plate. Firefly luciferase activity was measured using a Promega GloMax.

### HCV pseudoparticle (HCVpp) infectivity assay

To test HCVpp infectivity, Huh7.5 cells were seeded at 10,000 cells/well in a 96-well plate. The next day, cells were pre-treated with DMSO or Tg (100 nM) for 30 minutes and subsequently washed with 1X PBS. Treated cells were then infected with 50 µL/well of HCVpp and incubated for 3-4 hours. Following incubation, 100 µL of complete DMEM was added to each well. Cells were lysed 72 hours post infection using 0.1% Triton-X and combined with firefly luciferase reagent (Nanolight #318) and transferred to a white flat-bottom 96-well plate. Firefly luciferase activity was measured using a Promega GloMax.

### Lipoprotein secretion

Huh7.5 cells were seeded at 2 x 10^5^ cells/well in a 12-well plate overnight. The next day, cells were pre-treated with DMSO or 100 nM Tg for 30 minutes and subsequently mock or HCV-infected. Cellular supernatants were collected at 72 hpi, and 1% Triton-X was added to each sample to inactivate virus. ApoB secretion was measured using the Invitrogen Human ApoB ELISA kit (Thermo Fisher, #EH34RB) as per the manufacturer’s protocol. In parallel, the secretion of all other lipoproteins was assayed using the LEGENDplex Human Apolipoprotein multiplex bead-based assay (Biolegend, #740454) and measured using the Beckman Coulter Cytoflex S flow cytometer.

### Gaussia luciferase secretion

Huh7.5 cells were seeded at 2 x 10^5^ cells/well in a 12-well plate overnight. The next day, cells were transfected with 500 ng of pCMV-Gaussia luciferase (Gluc) using Fugene 6 for four hours in serum-free media. After 4 h, cells were treated with DMSO or Tg (100 nM) for 30 minutes, washed with 1X PBS and then replaced with complete DMEM. Four hours prior to the 24h harvest, cells were treated with 5 µg/mL brefeldin A (BFA; Biolegend #420601) for 30 minutes. Gluc activity was measured using coelenterazine (Nanolight #301). Briefly, 30 μL of supernatant was added to a white 96-well plate in triplicate corresponding to each well in the 12-well plate. The remaining supernatant was removed, and adhered cells were lysed with 1X passive lysis buffer (Nanolight #333) to assess intracellular Gluc activity. Cells were incubated in lysis buffer for 20 minutes, subsequently scraped, and 30 μL of lysate was transferred to a white 96-well plate (in triplicate). Luciferase expression was measured using a Promega Glomax.

### Cell viability assay

The viability of Huh7 or Huh7.5 cells in the presence of DMSO, Tg or IXA4 was assessed using the AlamarBlue reagent (Thermo Scientific), as per the manufacturer’s protocol.

### Statistical analysis

All statistical analysis was performed using GraphPad Prism for Mac version 10.1.1. We used a one way or two-way ANOVA and compared means regardless of row or column. We utilized a two-tailed unpaired Student’s t-test under the assumption of normality. Significance was calculated by prism as follows: *p<0.05, **p<0.01, ***p<0.0005, ****p<0.0001.

## ACKNOWLEDGEMENTS

This work was supported by the Canadian Institutes for Health Research (CCC; PTT-184008), the Natural Sciences and Engineering Research Council of Canada (CCC; RGPIN-2020–04277), and the Foundation for Innovation John R. Evans Leaders Fund (CCC). THT acknowledges support from a Vanier Canada Graduate Scholarship and a PhD studentship from the Canadian Network on Hepatitis C (CanHepC). CanHepC is funded by a joint initiative of the Canadian Institutes of Health Research (NHC-142832, NHE-174228, HPC-178912) and the Public Health Agency of Canada. SKS acknowledges an NSERC Canadian Graduate Scholarship-Doctoral. We thank Dr. Rodney Russell (Memorial University), Dr. Ralf Bartenschlager (Heidelberg University) and Apath LLC for providing HCV reagents. We are grateful to Dr. Abdullah Awadh (King Saud bin Abdulaziz University of Health Sciences) for advice on imaging lipid droplets, and to Dr. Craig McCormick (Dalhousie University, Canada) for providing reagents and insightful discussions.

## FIGURE LEGENDS

**Supplemental Table 1. RNA-Sequencing expression matrix.** Huh7.5 cells were treated in technical triplicate with DMSO or 100 nM Tg for 30 minutes, then washed and replaced with normal media. Cellular RNA was collected 48 h later. Samples were sent for Illumina 3′ end bulk RNA-sequencing.

**Supplemental Figure 1**. T**g upregulates genes involved in cholesterol biosynthesis and protein secretion**. Huh7.5 cells were treated with DMSO or 100 nM Tg for 30 minutes, then washed and replaced with normal media. Cellular RNA was harvested, 48 h post-treatment. Samples were sent for Illumina 3′ end bulk RNA-sequencing and pathway enrichment analysis was performed using the **(A)** Reactome Pathway or **(B)** Hallmark Pathway. Heatmap displays the Z-score of the most significantly upregulated genes in Tg treatment vs. DMSO control involved in **(A)** cholesterol biosynthesis or **(B)** protein secretion.

**Supplemental Figure 2**. C**e**ll **viability following treatment with DMSO, Tg or IXA4.** Huh7 cells were treated with DMSO and 100 nM Tg for 30 minutes or were treated with IXA4 (10 µM) and incubated for 72 h. Cell viability was measured using an Alamar blue assay.

