## Supplemental Figure 1 for "Modulation of lipid metabolism by thapsigargin inhibits hepatitis C virus infection by multiple mechanisms"

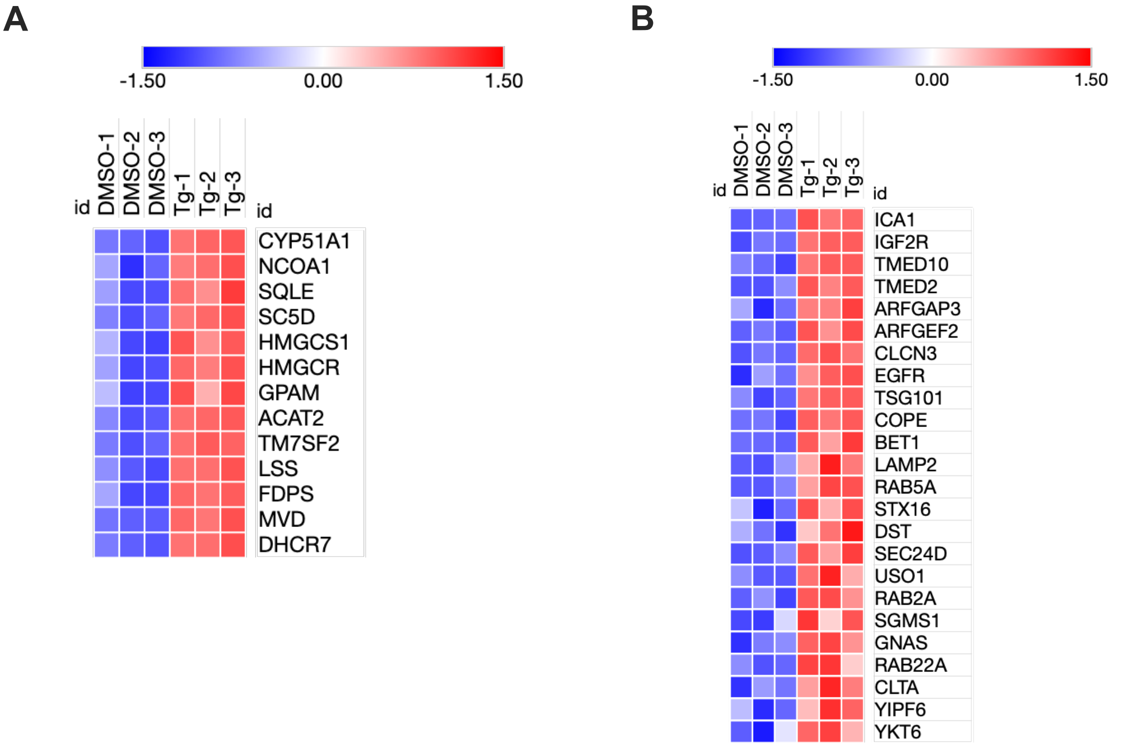
A
-1.50
0.00
1.50
id
DMSO-1
DMSO-2
DMSO-3
Tg-1
Tg-2
Tg-3
id
CYP51A1
NCOA1
SQLE
SC5D
HMGCS1
HMGCR
GPAM
ACAT2
TM7SF2
LSS
FDPS
MVD
DHCR7
B
-1.50
0.00
1.50
id
DMSO-1
DMSO-2
DMSO-3
Tg-1
Tg-2
Tg-3
id
ICA1
IGF2R
TMED10
TMED2
ARFGAP3
ARFGEF2
CLCN3
EGFR
TSG101
COPE
BET1
LAMP2
RAB5A
STX16
DST
SEC24D
USO1
RAB2A
SGMS1
GNAS
RAB22A
CLTA
YIPF6
YKT6
