## Supplementary figures and images for "Modulation of lipid metabolism by thapsigargin inhibits hepatitis C virus infection by multiple mechanisms"

### Supplemental Figure 2

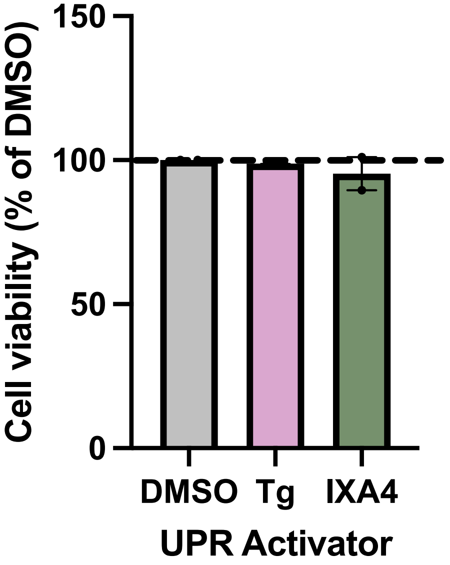

Cell viability (% of DMSO)
150
100
50
0
DMSO
Tg
IXA4
UPR Activator
